# Identification of novel Ebola virus inhibitors using biologically contained virus

**DOI:** 10.1101/2021.12.09.471933

**Authors:** Bert Vanmechelen, Joren Stroobants, Winston Chiu, Joost Schepers, Arnaud Marchand, Patrick Chaltin, Kurt Vermeire, Piet Maes

## Abstract

Despite recent advancements in the development of vaccines and monoclonal antibody therapies for Ebola virus disease, treatment options remain limited. Moreover, management and containment of Ebola virus outbreaks is often hindered by the remote nature of the locations in which the outbreaks originate. Small-molecule compounds offer the advantage of being relatively cheap and easy to produce, transport and store, making them an interesting modality for the development of novel therapeutics against Ebola virus disease. Furthermore, the repurposing of small-molecule compounds, previously developed for alternative applications, can aid in reducing the time needed to bring potential therapeutics from bench to bedside. For this purpose, the Medicines for Malaria Venture provides collections of previously developed small-molecule compounds for screening against other infectious diseases. In this study, we used biologically contained Ebola virus to screen over 4,200 small-molecule drugs and drug-like compounds provided by the Medicines for Malaria Venture (i.e., the Pandemic Response Box and the COVID Box) and the Centre for Drug Design and Discovery (CD3, KU Leuven, Belgium). In addition to confirming known Ebola virus inhibitors, illustrating the validity of our screening assays, we identified eight novel selective Ebola virus inhibitors. Although the inhibitory potential of these compounds remains to be validated in vivo, they represent interesting compounds for the study of potential interventions against Ebola virus disease and might serve as a basis for the development of new therapeutics.

## Introduction

Ebola virus (EBOV), previously known as Zaire Ebola virus, was first discovered in 1976 in the Democratic Republic of the Congo (previously called Zaire) and has since then caused several disease outbreaks, predominantly in central Africa [1,2]. Ebola virus disease (EVD) is a zoonotic hemorrhagic fever that, once introduced into humans, spreads from human-to-human via direct contact [3]. The incubation time varies from 2-21 days, after which symptoms develop suddenly, most frequently including fever, fatigue, headache, a sore throat and muscle pain, followed by vomiting, rash and diarrhea [4]. The average case fatality rate is 65%, although this varies strongly from outbreak to outbreak [5]. While vaccines have been developed that have been successfully used to limit the spread of EBOV outbreaks, treatment options for infected individuals are limited [6,7]. The current recommended treatment of Ebola patients is focused on early supportive care and symptomatic treatment, although a recent clinical trial found early-administered single-dose injections of two monoclonal antibodies, REGN-EB3 (Inmazeb) and MAb114 (Ebanga), to offer improvements in overall mortality [8]. A number of small-molecule compounds have also been evaluated as potential treatments for EVD, but none have proven efficacious in humans so far, potentially partly attributable to difficulties in establishing scientifically sound clinical trials in the field [9,10]. Even the nucleoside analogue remdesivir, which shows strong *in vitro* inhibition of EBOV and other negative-stranded RNA viruses, and which has been shown to efficiently protect non-human primates from EBOV challenge, failed to reduce mortality in the abovementioned trial [8,11,12]. However, it should be noted that only one dosing regimen was tested in this trial.

Even though the recent availability of efficacious vaccines and monoclonal antibodies have somewhat improved the outlook for future outbreak management, there remains a strong need for the discovery and development of more effective treatment options. However, because of the high risk these viruses pose, resulting in their classification as biosafety level 4 (BSL-4) agents, research with infectious virus is restricted to a limited number of BSL-4 facilities, hindering the rapid development of additional countermeasures [13]. To circumvent this need for BSL-4 laboratories, several virus alternatives have been developed that allow researchers to study EBOV in lower biosafety settings, including minigenome systems and virus-like particle systems [14]. Despite the limitations and drawbacks each of these systems has, their ability to be used in standard BSL-2 laboratories has resulted in these systems becoming commonly used tools for EBOV research and they have significantly boosted our knowledge and understanding of EBOV biology and outbreak management.

An alternative approach to studying EBOV in lower biosafety laboratories is by using ‘biologically contained EBOV’. In 2008, Peter Halfmann and colleagues showed that it is possible to confine EBOV to a specific cell line by removing one of the essential genes (VP30) from the virus genome and providing the missing protein in trans in the cell line of choice [15]. This results in the production of EBOV that is phenotypically near indistinguishable from wild-type virus, but which is safe to handle. Based on this principle, we created a biologically contained EBOV system in which the virus is confined to cell lines stably transduced with a lentiviral construct expressing EBOV VP30, while the VP30 gene in the virus genome is replaced by an eGFP reporter gene.

In this study, we used this system to screen two large repurposing compound libraries for their activity against EBOV. The first set consists of two compound libraries that are made freely available by the Medicines for Malaria Venture (MMV), the Pandemic Response Box and the COVID box, totaling 560 compounds. The Pandemic Response Box is a set of “*400 diverse drug-like molecules active against bacteria, viruses or fungi*” (MMV, The Pandemic Response Box, www.mmv.org/mmv-open/pandemic-response-box), while the COVID Box consists of 160 compounds *“with known or predicted activity against the coronavirus SARS-CoV-2”* (MMV, The COVID Box, www.mmv.org/mmv-open/covid-box)[16,17]. The second set is an in-house library provided by the Centre for Drug Design and Discovery (CD3, KU Leuven, Belgium), comprising more than 3,600 repurposing compounds. In addition to confirming known EBOV inhibitors, we identified several novel *in vitro* EBOV inhibitors, opening up new avenues for the development of novel EBOV therapeutics.

## Materials and methods

### Cell lines

Human embryonic kidney cells (HEK293FT; Thermo Fisher Scientific), Human hepatocellular carcinoma cells (Huh-7; Thermo Fisher Scientific) and African green monkey kidney cells (Vero E6; Vero C1008, ATCC) were passaged in DMEM (Thermo Fisher Scientific), supplemented with 10% FBS (Biowest) and 5% Penicilline-Strepomycin-Glutamine (Thermo Fisher Scientific). Additional supplements were 0.2% Amphotericin B (Thermo Fisher Scientific) and 2 µg/ml gentamicin (Thermo Fisher Scientific) for the Vero E6 cells, and 1% sodium bicarbonate (Thermo Fisher Scientific) and 1% NEAA (Thermo Fisher Scientific) for the Huh-7 cells. During assays, serum concentration was lowered to 2% for Vero E6 cells and 5% for Huh-7 cells.

### Plasmids

pCAGGS plasmids encoding the EBOV L, NP, VP30 and VP35 proteins, as well as a T7-3E-Luc-5E minigenome plasmid (all based on the Mayinga strain), were kindly provided by Prof. Stephan Becker. A plasmid encoding an eGFP-containing EBOV antigenome was generated through assembly of fragments using the NEBuilder HiFi DNA assembly cloning kit (New England Biolabs (NEB)). The vector backbone, T7 promoter, virus leader, virus trailer, HdVRz and T7 terminator sequences were derived from the T7-3E-Luc-5E vector. Fragments covering the NP and L genes were derived from the corresponding pCAGGS helper plasmids. The rest of the antigenome, from the intergenic region in front of the VP35 gene to the intergenic region behind the VP24 gene, with an eGFP gene replacing the VP30 coding region, was synthesized in a pUC57 vector by GenScript Biotech based on EBOV strain Mayinga (GenBank: AF272001). This fragment was digested with NotI-HF and SmaI (NEB), while all other fragments were amplified by PCR using the Q5 HotStart High-Fidelity 2X master mix (NEB). All plasmids were sequence-verified with Sanger sequencing (Macrogen Europe, Amsterdam, The Netherlands) before use.

### Lentiviral constructs

Cell lines expressing VP30 were made by lentiviral transduction. Using the NEBuilder HiFi DNA assembly cloning kit (NEB), EBOV VP30 was inserted into a pLenti6.3 vector (Thermo Fisher Scientific), in which the CMV promoter was replaced by an SFFV promoter derived from a pHR-SFFV-dCas9-BFP-KRAB vector (www.addgene.org, Cat. #46911). An internal ribosomal entry site (IRES) cassette was inserted between the VP30 gene and the blasticidin resistance marker by restriction enzyme digestion of the vector with SpeI-HF and SalI-HF (NEB), and digestion of a pEF1a-IRES vector (www.takarabio.com, Cat. # 631970) with NheI-HF and SalI-HF (NEB). Fragments were ligated with the Quick Ligation kit (NEB).

### Lentiviral transduction

For lentivirus production, 50-70% confluent HEK293FT cells in T-25 flasks were transfected with Lipofectamine LTX & PLUS Reagent (Thermo Fisher Scientific). LTX solution and transfection mixes containing 3 µg of lentiviral EBOV vector, 5.83 µg of psPAX2 vector, 3.17 µg of pMD2.G vector and 12 µL PLUS reagent were prepared in serum-free Opti-MEM (Thermo Fisher Scientific). Following a five-minute incubation at room temperature, solutions were mixed and incubated for an additional 20 minutes. Cell medium was replaced by 5 mL of fresh medium, after which transfection complexes were added, followed by a 21-hour incubation at 37°C. Next, sodium butyrate (10 mM) was added and cells were incubated for an additional 3 hours, after which the medium was replaced with 5 mL of fresh medium. Virus-containing supernatants were harvested into 15 mL conical tubes 24 hours after sodium butyrate addition and centrifuged at 2000g for 15 minutes at 4°C to pellet cell debris. Transduction of cell lines with the harvested lentivirus was done according to the ViraPower HiPerform T-Rex Gateway Expression System (Thermo Fisher Scientific) manufacturer’s protocol. Six µg/ml Polybrene (Sigma-Aldrich, Saint-Louis, MO, USA) was used to increase transduction efficiency. Following transduction, cell medium was supplemented with 10 µg/ml blasticidin (InvivoGen) during passaging.

### EBOV rescue

Huh-7 cells transduced with EBOV VP30 (Huh-7-EBOV-VP30) were seeded in a 6-well plate (300.000 cells/well). Following overnight incubation, the cells were transfected with 1000 ng EBOV antigenome, 1000 ng T7 polymerase, 1000 ng pCAGGS-EBOV-NP, 2000 ng pCAGGS-EBOV-L, 500 ng pCAGGS-EBOV-VP35 and 500 ng pCAGGS-EBOV-VP30, using 3:1 TransIT-LT1 Transfection Reagent (Mirus Bio). Twenty-four hours later, the medium was replaced by fresh medium. Six days post-transfection, cells were trypsinized and mixed with fresh Huh-7-EBOV-VP30 cells in a T-25 flask. After three days, supernatant from flasks showing widespread eGFP expression was collected and used to infect Vero E6-EBOV-VP30 cells seeded one day prior in a T-25 flask. After six days, the supernatant was used to infect additional T-75 flasks of Vero E6-EBOV-VP30 cells, from which, after seven days, the supernatant was collected, centrifuged at 17.000g for three minutes and subsequently aliquoted and stored at -80°C.

### RNA extraction and nanopore sequencing

RNA was extracted from 100 µl of virus stock using a KingFisher Flex (Thermo Fisher Scientific) in combination with the MagMax Viral Pathogen kit II (Thermo Fisher Scientific), according to the manufacturer’s instructions. RNA was converted to cDNA and amplified by Sequence-Independent Single Primer Amplification as described by Greninger et al. [18]. The resulting cDNA was prepared for nanopore sequencing using the SQK-LSK110 kit (Oxford Nanopore Technologies (ONT), Oxford, UK) with the EXP-NBD114 barcoding expansion (ONT). The resulting library was loaded on a R9.4.1 flow cell and run on a GridION. Basecalling and barcode demultiplexing was done using the ont-guppy-for-gridion v4.2.3. The resulting reads were mapped against the plasmid design used for generation of the antigenome construct using Minimap2 v2.17-r941, followed by Medaka v1.0.1 for consensus polishing and variant calling [19].

### Virus titration

Vero E6-EBOV-VP30 cells were seeded in 6-well plates. Once confluent, cell medium was removed and 200 µl virus dilution was added to each well. A ten-fold dilution series, covering ten dilutions (1×10^-1 -1×10^-10) was used, with duplicate repeats for each concentration. Plates were kept in an incubator (37°C, 5% CO2), gently swirling the plates every 15 minutes. After one hour, 3 ml freshly prepared agarose-medium was added to each well. Agarose-medium was made by autoclaving a 17.6 µg/ml SeaKem ME agarose (Lonza, Basel, Switzerland) dilution and heating it to 65°C. Once heated, the agarose was added to preheated (37°C) 2X Basal Medium Eagle without Earle’s salts (Thermo Fisher Scientific), supplemented with 10% FBS (Biowest), 200 mM L-glutamine, 1% NEAA, 1% Penicillin-Streptomycin, 1% Gentamicin and 0.2% Fungizone (all Thermo Fisher Scientific), in a 1:2 ratio. After cooling down to room temperature, plates were moved to an incubator for five days. Read-out was performed by counting the amount of eGFP+-cell clusters.

### Antiviral screening assay

Compounds, spotted in 96-well plates at 2 or 10 mM, were gifted to us by MMV and CD3. Intermediary compound dilutions were made in complete cell medium directly before adding the compound to 96-well plates (CELLSTAR, Greiner-Bio, Vilvoorde, Belgium) in which Vero E6-EBOV-VP30 cells had been seeded one day prior at 20,000 cells/well. For the MMV compound set, a dilution series of four concentrations was tested for each compound, starting at 50 µM and diluting four-fold each time, allowing twenty-two compounds to be tested per plate. For the CD3 set, two concentrations (1 and 10 µM) were tested for each compound on separate plates. Following compound addition, 200 plaque forming units (PFU) virus dilution was added to each well. Medium without virus was added to the negative controls. Six days post-infection, cell medium was replaced by fresh medium supplemented with 5 µM Hoechst 33342 nucleic acid stain (Thermo Fisher Scientific) as a background stain for high-content imaging analysis. Imaging and image analysis was done using an Arrayscan XTI (Thermo Fisher Scientific) and a custom Cellomics SpotDetector BioApplication protocol, as described previously [20]. Further data analysis was done using Genedata Screener V17.05-Standard. GraphPad Prism v8.2.0 was used for graph plotting.

### Hit confirmation

To confirm compound activity observed in the initial screening assays, additional compound was acquired. For the MMV compounds, fresh DMSO stocks were prepared from powder provided by Evotec (Hamburg, Germany), while the CD3 compounds were provided as DMSO solutions. Hit confirmation using these fresh stocks was done by testing each compound in triplicate in Vero E6-EBOV-VP30 cells over a two-fold dilution series of nine dilutions, starting at 100 µM. Cell and virus quantities were identical to the ones used in the screening assay and plate handling procedures and data read-out were performed as described above. In addition to Vero E6-EBOV-VP30 cells, a subset of compounds was also tested in Huh-7-EBOV-VP30 cells. In this cell line, a two-fold dilution series of eight dilutions, starting at 50 µM, was used. To allow adequate high-content imaging, Huh-7-EBOV-VP30 cells were seeded at 10,000 cells/well and infected with 0.2 PFU EBOV-ΔVP30-eGFP per cell. Assay read-out was performed four days post-infection. Other plate handling and data processing procedures were performed as described above.

## Results

### Rescue and characterization of biologically contained EBOV

To set up a screening platform for EBOV inhibitors using infectious virus without requiring access to a BSL-4 facility, we created a biologically contained EBOV system similar to the one described by Halfmann et al. [15]. EBOV VP30-expressing cell lines were generated by lentiviral transduction of Vero E6 and Huh-7 cells with a lentiviral vector in which the VP30 gene was coupled to a blasticidin resistance marker by means of an IRES. Rescue of biologically contained EBOV was then attempted by transfecting the selected Huh-7-EBOV-VP30 cells with a VP30-deficient EBOV antigenome containing an eGFP gene, under the control of a T7 polymerase promoter. Six days post-transfection with the antigenome construct and the necessary support plasmids, extensive cell death was observed in all wells. Only in one of the Huh-7-EBOV-VP30 wells, one cluster (∼30 cells) of green cells could be observed. These cells were collected and mixed with fresh Huh-7-EBOV-VP30 cells. After 72 hours, the medium of these cells was collected and used to infect Vero E6-EBOV-VP30 cells, resulting in widespread eGFP-expression six days post-infection. Supernatant from these cells was used to infect additional flasks of Vero E6-EBOV-VP30 cells, to generate a large stock of EBOV-ΔVP30-eGFP virus. Nanopore sequencing of this stock revealed five acquired mutations compared to the construct from which it was derived: two non-synonymous mutations in the NP gene (T928C -> S524F, G2551A -> F648L) and three mutations in the L gene (G14038A -> A820T, G14187A (silent), G18138A -> M2186I).

To assess the usability of EBOV-ΔVP30-eGFP for compound screening, we determined the minimum infectious dose needed to obtain widespread eGFP expression in Vero E6-EBOV-VP30 cells seeded in 96-well plates. A loading dose of 200 PFU/well, corresponding to 0.01 PFU/cell, was found to yield >85% eGFP-positive cells six days post-infection in all replicates, while lower doses failed to uniformly infect all cells or replicates (Figure 1A). To evaluate the growth kinetics of all virus isolates in greater detail, virus growth was observed daily over a period of six days, using the minimal titer required for optimal growth (0.01 PFU/cell), as well as a 10-fold higher dose (0.1 PFU/cell). In both conditions, initial eGFP expression could be observed already after one day, increasing rapidly until day 4-6, with higher titers yielding faster virus propagation, as evidenced by uniform eGFP expression of the infected cells (Figure 1B). When using non-transduced Vero E6 cells, no eGFP expression was observed, regardless of the used loading dose, confirming the confinement of EBOV-ΔVP30-eGFP to the VP30-transduced Vero E6 cells. To further validate this confinement, the virus was passaged an additional six times on Vero E6-EBOV-VP30 cells. Supernatans from each passage was used to infect fresh transduced and untransduced Vero E6 cells. For each passage, >95% eGFP expression could be observed in all replicates of the transduced cells six days post-infection, while no eGFP expression was ever observed in untransduced cells.

**Figure 1:**
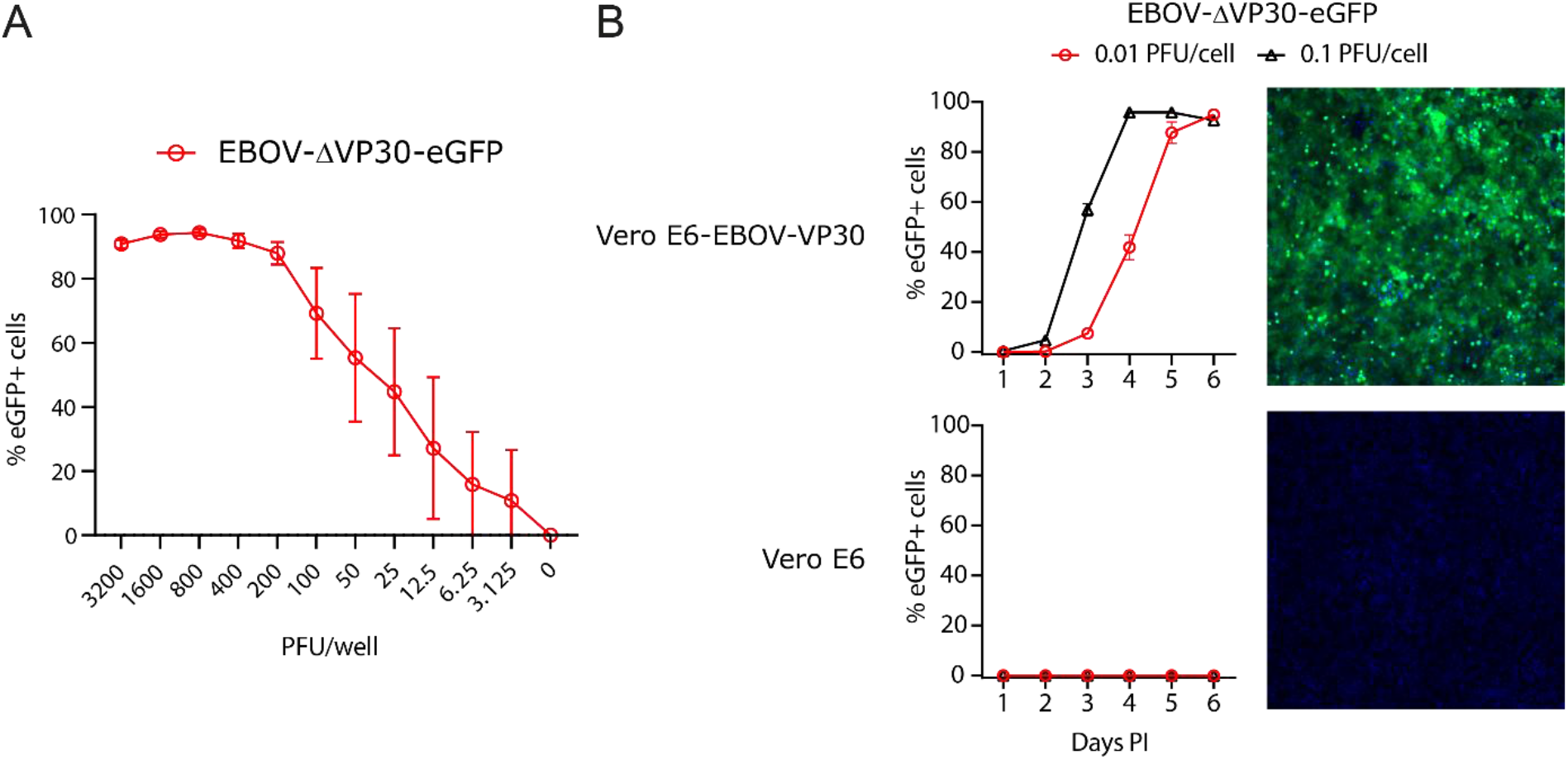
EBOV-ΔVP30-eGFP characterization. **(A)** Minimal virus titer required for homogenous infection in 96-well plates. Vero E6-EBOV-VP30 cells were infected with EBOV-ΔVP30-eGFP and the fraction of eGFP-positive cells was determined by high-content imaging six days post-infection. Different viral titers tested are expressed in plaque forming units (PFU)/well. Eight replicates across two separate plates were performed per condition. Error bars denote standard deviation. **(B)** Growth kinetics of EBOV-ΔVP30-eGFP in Vero E6-EBOV-VP30 and regular Vero E6 cells. The left graph shows the fraction of eGFP-positive cells, measured by high-content imaging. The right column shows a representative image of a well infected with 0.1 PFU/well, five days post-infection. Green cells express eGFP and all cells are background stained with Hoechst 33342 (blue). PI = post-infection, PFU = plaque forming units. At least three replicates are included for each condition. Error bars denote standard deviation.

### Compound screening

Once validated for safety, our EBOV-ΔVP30-eGFP assay was subsequently used to screen >4,000 compounds for their potential as EBOV inhibitors. 560 compounds were obtained from MMV as part of the Pandemic Response Box and the Covid Box, while an additional 3,681 compounds were provided by CD3. High-content imaging was used to simultaneously assess antiviral activity and toxicity. For practical reasons, the initial screens of the MMV and CD3 libraries were performed separately, using two different plate layouts (Figure 2). Both the Pandemic Response Box (400 compounds) and the COVID Box (160 compounds) were initially screened over a 1:4 dilution range of four concentrations, starting at 50 µM, or 10 µM if the compounds were delivered at a lower starting concentration. This initial screen had an average Z’-factor of 0.89, calculated by comparing the means and deviations of the negative and positive controls of each plate using the following formula: Z’ = 1-[(3_σ_C_+_ + 3_σ_C_-_)/(_µ_C_+_ - _µ_C_-_)] [21]. Sixteen compounds (2.9%) showed a decrease in relative virus growth of >40% whilst maintaining >40% cell viability in at least one concentration tested. These compounds were retested in duplicate over a wider concentration range to rule out false positives. In this confirmation assay, two compounds failed to show significant virus inhibition and three compounds were insufficiently selective (estimated selectivity index (SI) <3). These five compounds were excluded from further analysis. The CD3 compound library consisted of 3,681 compounds selected from marketed/withdrawn drugs, compounds currently in clinical trials and annotated bioactive molecules. All compounds were pre-spotted in 96-well plates at a stock concentration of 10 mM. Initial screening of these compounds was performed at working concentrations of 1 and 10 µM. The global Z’-factor for this screen was 0.87. Sixty-one compounds (1.7%) that showed more than 80% inhibition of eGFP-expression compared to the control while simultaneously showing less than 20% reduction in cell number were selected as preliminary hits (Figure 3). Comparably to the preliminary hits of the MMV screen, these 61 compounds were retested in duplicate over a wider concentration range to rule out false positives. Twenty-one compounds that showed an SI >5 and an IC_50_ <15 µM, in this confirmation assay, were retained for further analysis.

**Figure 2:**
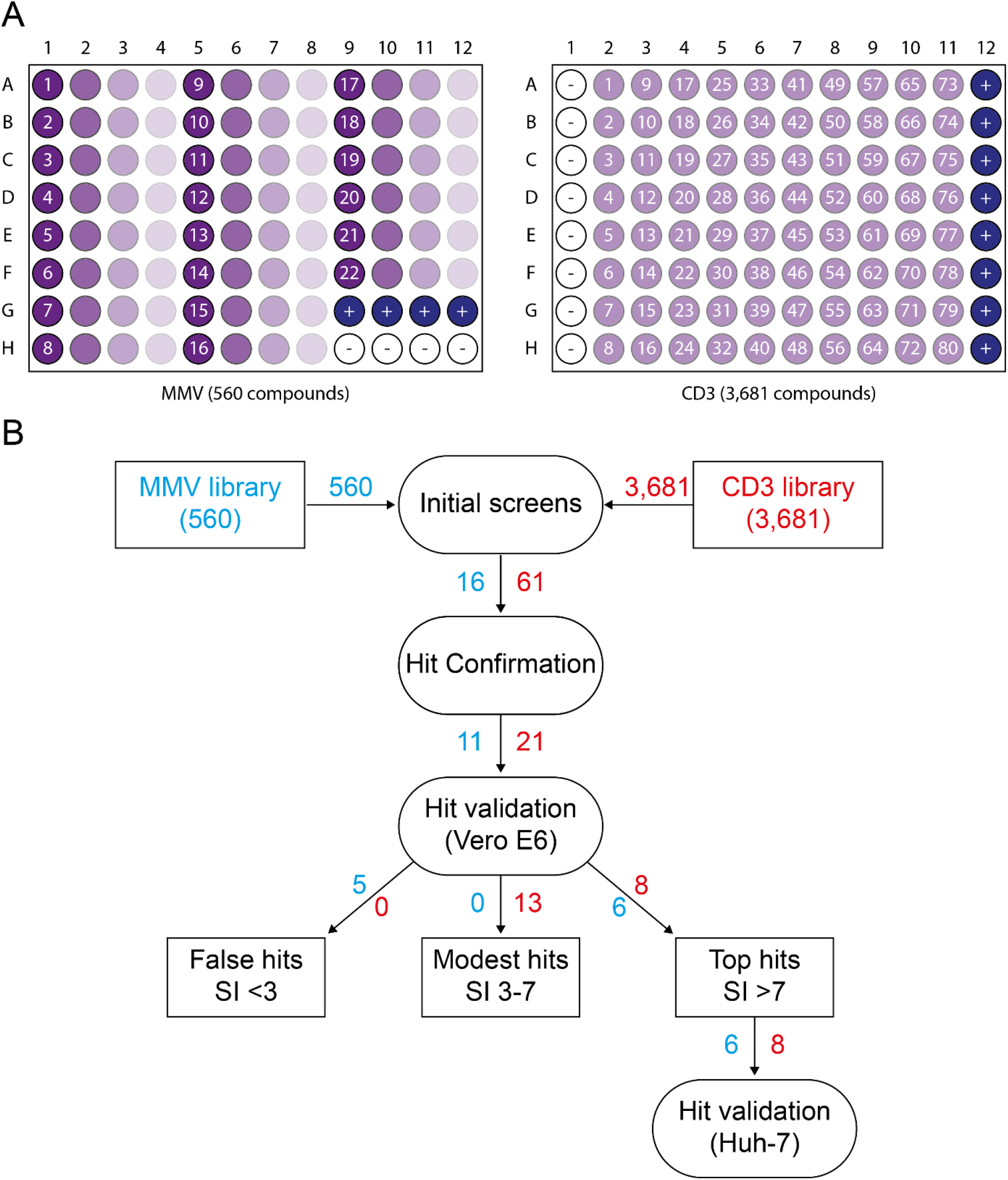
Screening assays layout. **(A)** Schematic representation of the assay layout used for the two different compound libraries. A single 1:4 dilution series was tested for each of the MMV compounds, while each CD3 compound was assayed twice, once at 1 µM and once at 10 µM. + = positive control (virus, no compound), - = negative control (no virus, no compound). **(B)** Schematic overview of the different assays performed. Numbers next to the arrows indicate the compounds continuing to the next step.

**Figure 3:**
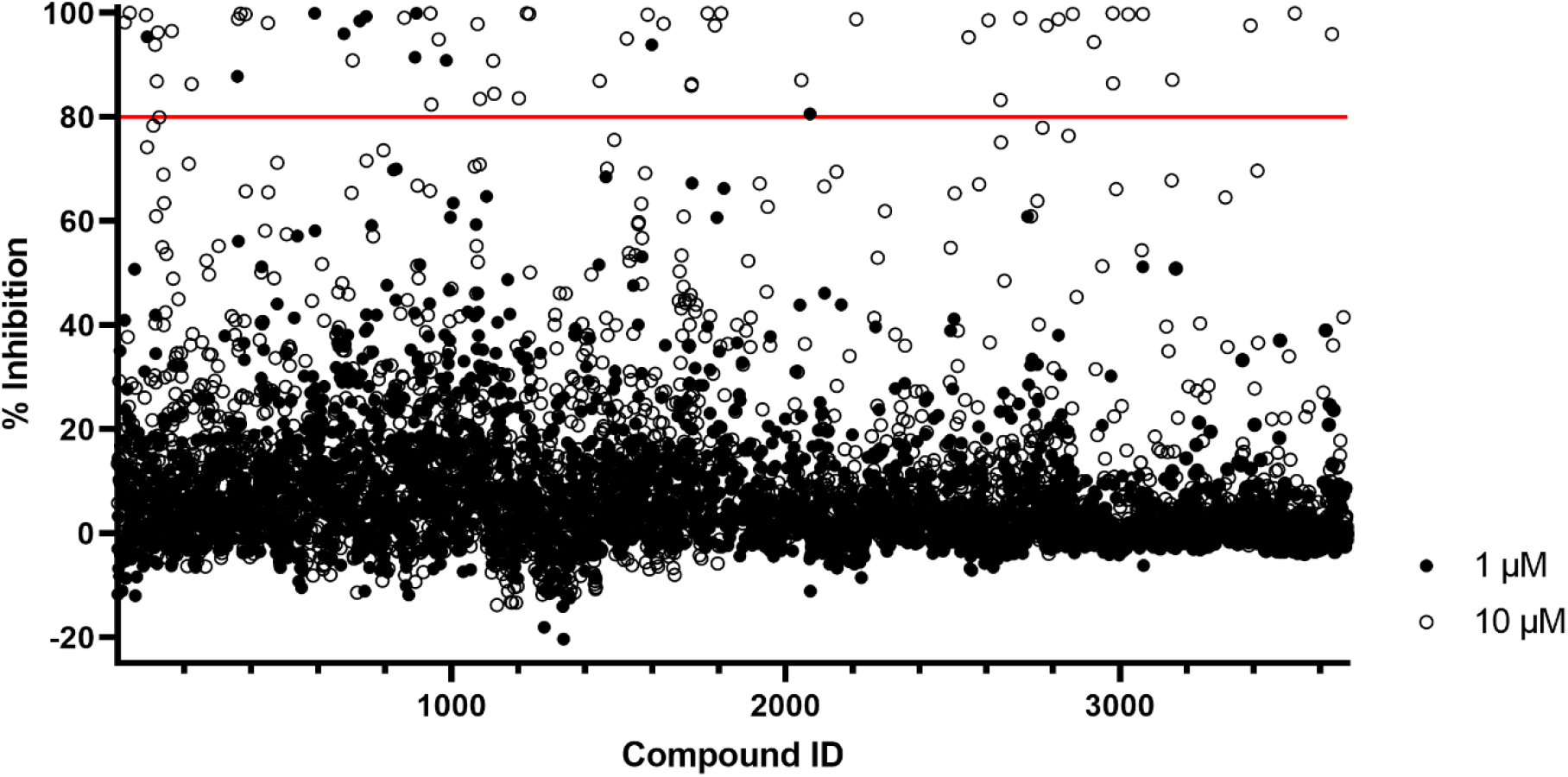
CD3 library preliminary hit selection. Overview of the initial screening results for all 3,681 CD3 compounds. Each compound was tested at 1 and 10 µM. Only compounds that showed less than 20% reduction in cell survival are shown. Shown on the Y-axis is the relative inhibition of eGFP-expression compared to the non-compound control. Compounds that reduced eGFP-expression by >80% in either concentration were selected as preliminary hits.

### Hit validation

New aliquots of the antivirals in the form of compound powder (MMV) or DMSO solution (CD3) were acquired to confirm the activity and selectivity of the eleven MMV and twenty-one CD3 compounds that had demonstrated selective inhibition of EBOV replication. Triplicate testing of the new compound stocks over a range of nine concentrations was used to accurately determine IC_50_, CC_50_ and SI values for each compound (Table 1). Twenty-seven of the thirty-two preliminary hits were confirmed to inhibit EBOV-ΔVP30-eGFP replication, although several compounds seemed to be only moderately selective. The confirmed hits include two duplicates, itraconazole and retapamulin, which were present in both the MMV and CD3 compound libraries. Four of the five MMV compounds that failed to be confirmed (pimozide, apremilast, dabrafenib and fluconazole) were also present in the CD3 compound library but had not been picked up as hits in the CD3 screen, confirming their lack of potency. Comparably, one of the weaker hits of the CD3 compounds (benztropine) was also present in the MMV library but had failed to meet the criteria for initial hit selection. The fourteen compounds that showed the highest SI (all >7) in Vero E6 cells, including the duplicates of itraconazole and retapamulin, were retested using Huh-7-EBOV-VP30 cells (Table 1; Figure 4). The selective inhibition (SI >3) of EBOV-ΔVP30-eGFP replication was confirmed in Huh-7 cells for most compounds, with the exception of itraconazole, MMV1782214 and doramapimod. Interestingly, dalbavancin and benzyloxycarbonyl-phenylalanine-alanine-fluoromethylketone (z-FA-FMK) were notably more potent in Huh-7 cells, without apparent toxicity. The selective activity of apilimod and diphyllin in Huh-7 cells could not be accurately assessed because both their IC_50_ and CC_50_ values fell (almost) outside the tested concentration range.

**Table 1:** Activity and toxicity of hit compounds.

| Compound ID | Compound name | Vero E6 |  |  | Huh-7 |  |  |
| --- | --- | --- | --- | --- | --- | --- | --- |
|  |  | IC<sub>50</sub> (μM) | CC<sub>50</sub> (μM) | SI | IC<sub>50</sub> (μM) | CC<sub>50</sub> (μM) | SI |
| MMV637528<sup>¶</sup> | Itraconazole | 0.51 | >100 | >196 | 1.26 | 1.72 | 1.40 |
| MMV1804187 | Apilimod | <0.08 | 10.0 | >129 | <0.39 | <0.39 | - |
| MMV1803859 | Remdesivir | 0.67 | 55.9 | 83.1 | <0.39 | 2.00 | >5.12 |
| CD1260<sup>¶</sup> | Itraconazole | 1.51 | >100 | >66.4 | 2.58 | 2.49 | 0.96 |
| CD3051 | Z-FA-FMK | 2.30 | >100 | >43.5 | <0.39 | >50 | >128 |
| CD2833 | Evans blue | 3.07 | >100 | >32.6 | 9.92 | >50 | >5.04 |
| MMV1782214 | CHEMBL93139* | 1.30 | 36.4 | 27.9 | 3.61 | 9.15 | 2.54 |
| MMV1633674<sup>¶</sup> | Retapamulin | 3.72 | 65.0 | 17.5 | 0.48 | 8.16 | 16.9 |
| CD2986 | UNC1999 | 5.90 | 96.0 | 16.3 | 3.58 | 24.7 | 6.91 |
| CD2794<sup>¶</sup> | Retapamulin | 7.27 | 83.9 | 11.5 | 0.79 | 11.1 | 14.1 |
| CD2842 | Dalbavancin | 8.03 | 91.2 | 11.4 | 2.12 | >50 | >23.6 |
| CD3372 | Benproperine | 7.04 | 70.5 | 10.0 | 1.56 | 13.0 | 8.31 |
| CD2035 | Doramapimod | 10.8 | >100 | >9.23 | 8.84 | 10.8 | 1.22 |
| MMV102270 | Diphyllin | 0.42 | 3.09 | 7.44 | <0.39 | 0.43 | >1.10 |
| CD1581 | Oxelaidin | 12.8 | 85.8 | 6.71 | - | - | - |
| CD1056 | Chlorophyllin | 7.47 | 50.0 | 6.70 | - | - | - |
| CD3648 | Buclizine | 6.09 | 40.5 | 6.65 | - | - | - |
| CD3561 | Micafungin | 16.9 | >100 | >5.91 | - | - | - |
| CD2486 | GW-5074 | 4.95 | 23.0 | 4.64 | - | - | - |
| CD2111 | Pitavastatin | 0.88 | 4.07 | 4.61 | - | - | - |
| CD0329 | Tamoxifen | 4.11 | 18.7 | 4.56 | - | - | - |
| CD2889 | Propiverine | 10.1 | 43.8 | 4.36 | - | - | - |
| CD0888 | Benztropine | 7.90 | 32.9 | 4.16 | - | - | - |
| CD0842 | Carvedilol | 7.41 | 29.9 | 4.04 | - | - | - |
| CD3124 | Pilaralisib | 6.58 | 24.2 | 3.68 | - | - | - |
| CD1724 | Hydroquinidine | 27.0 | 82.1 | 3.04 | - | - | - |
| CD2145 | Fluvastatin | 1.83 | 5.56 | 3.03 | - | - | - |
| MMV002137 | Pimozide | 2.77 | 7.83 | 2.82 | - | - | - |
| MMV1804411 | (RS)-PPCC | 58.3 | >100 | >1.71 | - | - | - |
| MMV1804482 | Apremilast | 82.3 | >100 | >1.22 | - | - | - |
| MMV1803334 | Dabrafenib | 33.6 | 27.4 | 0.81 | - | - | - |
| MMV002337 | Fluconazole | >100 | >100 | - | - | - | - |

<sup>¶</sup>Duplicate compound present in both the MMV and CD3 libraries
\*ChEMBL identifier

**Figure 4:**
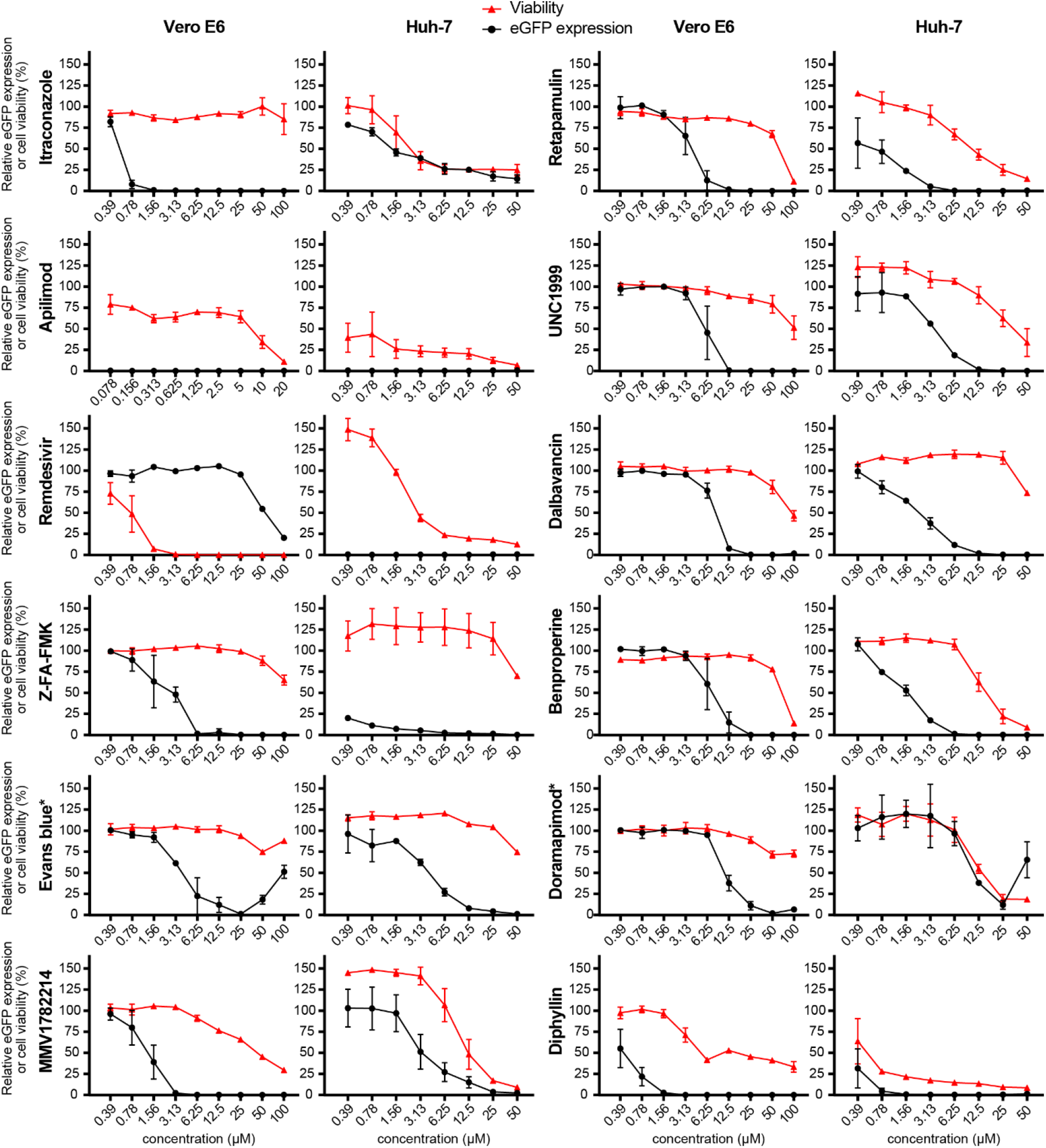
Activity and toxicity profiles of the twelve compounds that showed the highest selectivity index in Vero E6 cells. All graphs indicate relative cell survival (red triangles) and relative eGFP expression (black circles) compared to the positive control (no compound). Left graphs indicate the results obtained using Vero E6-EBOV-VP30 cells, while right graphs represent the results from Huh-7-EBOV-VP30 cells. For itraconazole (MMV637528) and retapamulin (MMV1633674), only the values obtained using the most recent stocks are shown. *The increase in eGFP expression at higher concentrations for these compounds is not caused by virus growth but by excess compound causing light scattering within the eGFP spectrum. Error bars indicate standard deviation. Data from three (Vero E6) or six (Huh-7) replicates.

## Discussion

Despite recent advancements in the search for EBOV therapeutics, no small-molecule compounds are licensed to treat EBOV infections [9]. However, small-molecule compounds are generally easy and cheap to produce, transport and store, making them interesting candidates for the treatment of patients, especially in remote locations [22,23]. Additionally, because many small-compound libraries have already been developed for a variety of applications, the repurposing of existing compounds forms an interesting research avenue for the rapid identification and implementation of potential antivirals. In this study, we optimized a biologically contained EBOV assay and used it to screen the MMV Pandemic Response box and COVID Box, two such libraries of small-molecule compounds with drug-like characteristics that have been independently developed for the antimicrobial treatment of various infections [16,17]. Additionally, we screened a large in-house repurposing collection, provided by CD3. In total, 4,241 compounds from these three libraries were tested for their anti-EBOV potential. Thirty unique compounds were retained after the initial screens, twelve of which were ultimately found to profoundly inhibit EBOV-ΔVP30-eGFP replication with an SI >7 in at least one of the cell lines tested.

Four of the most active compounds, remdesivir, apilimod, diphyllin and dalbavancin, were previously identified as EBOV inhibitors [24,25]. Remdesivir is an adenosine analogue monophosphoramidate prodrug that is known to inhibit the polymerase activity of many mononegaviruses, including members of the families *Pneumoviridae, Paramyxoviridae* and *Filoviridae* [11]. It is also a known inhibitor of coronaviruses, including severe acute respiratory syndrome coronavirus 2 (SARS-CoV-2) [26,27]. In addition to showing excellent *in vitro* activity against filoviruses, remdesivir has been reported to be an effective post-exposure treatment for EBOV infection *in vivo*, as it was found to ameliorate disease symptoms and improve survival rates in a non-human primate model [12]. However, as mentioned above, despite showing excellent *in vitro* and *in vivo* anti-EBOV potential, remdesivir failed to improve the survival rates of EVD patients during a clinical trial carried out during the 2018-2020 EBOV outbreak in the Democratic Republic of the Congo [8]. Furthermore, also for the treatment of COVID-19, caused by SARS-CoV-2, the benefit of remdesivir is heavily contested [28-31]. Unlike remdesivir, the potential of apilimod as an EBOV inhibitor has not yet been evaluated *in vivo*. Conversely, *in vitro*, apilimod has been shown to potently inhibit EBOV replication in Vero E6, Huh-7 and primary human macrophage cells [24]. Apilimod was first identified as an inhibitor of Toll-like receptor-mediate interleuking-12/-23 signaling and has been evaluated as a potential anti-inflammatory drug for the treatment of Crohn’s disease, rheumatoid arthritis and psoriasis, albeit without significant clinical success [32-34]. Later research showed that apilimod works by inhibiting phosphatidylinositol-3-phosphate 5-kinase (PIKfyve), a lipid kinase involved in maintaining endosome morphology and ensuring endosome maturation [35]. By inhibiting PIKfyve and preventing endosome maturations, apilimod is believed to block EBOV entry, as endosome maturation is a crucial process needed to allow the EBOV GP to be cleaved by cathepsins L and B, exposing the GP receptor-binding domain and enabling binding of the EBOV entry receptor NPC1 [36,37]. Because it is well tolerated in humans and targets a rather unique part of the virus life cycle, apilimod is an interesting candidate to be part of combination therapies for the treatment of EBOV, but future research will first need to confirm its *in vivo* efficacy and clinical benefit. Comparable to apilimod, diphyllin and its derivatives interfere with EBOV entry by preventing endosome maturation [38]. Diphyllin is an inhibitor of vacuolar-type ATPase (V-ATPase), which hydrolyses adenosine triphosphate and simultaneously transports protons across cellular membranes, resulting in endosome acidification [39]. Lastly, dalbavancin, a glycopeptide antibiotic primarily used for the treatment of skin and soft tissue infections, is known to inhibit cellular entry of many different viruses, including echovirus 1, severe acute respiratory syndrome coronavirus, Middle East respiratory syndrome–related coronavirus and EBOV [40-42]. Unlike apilimod and diphyllin, which target endosome maturation, dalbavancin prevents virus entry by direct inhibition of cathepsin L [42]. In the case of EBOV, this results in the GP being kept in its pre-cleaved state, rendering it unable to bind NPC1 [43].

In the MMV compound library, three additional compounds, itraconazole, retapamulin and the yet-unnamed MMV1782214 (CHEMBL93139), were found to selectively inhibit EBOV-ΔVP30-eGFP replication. The former two compounds were also present and identified as hits in the CD3 library. Itraconazole, the top hit in both compound libraries, is a triazole derivate that works as a broad-spectrum antifungal agent [44]. Although it can cause mild gastrointestinal disturbances, cardiotoxicity and hepatotoxicity, itraconazole is generally well tolerated and it is used for the treatment of systemic (histoplasmosis, aspergillosis, blastomycosis) and superficial (onychomycosis) fungal infections [45]. In Vero E6 cells, itraconazole shows anti-EBOV activity in the sub-micromolar range without apparent toxicity. Conversely, in a hepatocyte cell line (Huh-7), itraconazole shows significant toxicity and no selective inhibition of EBOV replication. In addition to its antifungal properties, recent research has identified itraconazole as a potential cancer treatment and as an inhibitor of several viruses, including influenza virus, enteroviruses and coronaviruses [46-50]. The mechanisms for these newly discovered functions appear unrelated to its antifungal properties. Although not fully understood yet, itraconazole affects angiogenesis through indirect inhibition of the mechanistic Target of Rapamycin (mTOR), an important oncogene that regulates cell growth and proliferation [51]. One of the mechanisms behind the inhibition of mTOR activity by itraconazole appears to be dysregulation of cholesterol trafficking, which results in endosomal cholesterol accumulation [51,52]. This disturbance of cellular cholesterol homeostasis in itself is believed to contribute to the antiviral properties of itraconazole, by preventing virus escape from the endosome and simultaneously interfering with virus egress [49]. Interestingly, the mechanism behind impaired cholesterol trafficking has been shown to be a direct interaction between the cholesterol transporter NPC1 and itraconazole [51,53]. NPC1 is known to be an indispensable entry receptor for EBOV and the interaction between NPC1 and the EBOV GP in mature endolysosomes is necessary for the release of the virion contents into the cellular cytoplasm [54,55]. Possibly, the interaction between itraconazole and the EBOV entry receptor NPC1 further potentiates the antiviral activity of this compound in filovirus infections, although further research is necessary to fully elucidate the interaction of itraconazole with both host and viral factors. The second hit compound that was present in both libraries is retapamulin, a derivative of pleuromutilin approved for use in humans [56]. This compound showed comparable selectivity in both Vero E6 and Huh-7 cells, although it was roughly one order of magnitude more potent in the latter. Unlike the aforementioned compounds, retapamulin has not yet been reported to possess antiviral activity and its sole known function is the inhibition of the bacterial ribosome complex [57]. While retapamulin is only used for topical application, other pleuromutilins can be used systemically and might be of use for EVD treatment, although their antiviral mechanism of action would first need to be elucidated [58]. Lastly, MMV1782214 (CHEMBL93139) showed excellent selectivity in Vero E6 cells but less so in Huh-7 cells. This compound is a 1,3,4-trisubstituted pyrrolidine derivative that was developed as a C-C chemokine receptor type 5 (CCR5) antagonist to be used for the treatment of human immunodeficiency virus infections [59]. Other pyrrolidine derivatives have recently been shown to inhibit EBOV replication, although the mechanism through which this inhibition is achieved remains to be determined [60].

In the CD3 library, more than twenty compounds displayed selective anti-EBOV activity, although for most compounds this selectivity was modest (SI 3-7). Aside from the aforementioned known EBOV inhibitors and compounds also present in the MMV library, five compounds were found to show strong selectivity towards virus inhibition: z-FA-FMK, Evans blue, UNC1999, benproperine and doramapimod. Z-FA-FMK is a potent inhibitor of cysteine proteases, including cathepsin B and L [61]. As mentioned above, these cathepsins are needed to cleave the EBOV GP before it can interact with NPC1. In both Vero E6 and Huh-7 cells, z-FA-FMK showed only limited toxicity while inhibiting EBOV-ΔVP30-eGFP in the low-micromolar or even sub-micromolar range, presumably by preventing virus entry, making it an interesting putative EBOV therapeutic. However, because of z-FA-FMK’s broad and potent inhibitory effect on cysteine proteases and its known function as an immunosuppressant that can interfere with T-cell proliferation, detailed *in vivo* validation of its safety and clinical benefit would be needed before it could be considered for use in humans [62]. For Evans blue, UNC1999 and benproperine, the mechanism through which they might inhibit EBOV replication is less clear. Evans blue or T-1824 is an azo dye that is known for its dark blue color and high affinity for albumin, and it is primarily used to stain cells or tissues in a laboratory setting [63]. However, it is also known to bind several glutamate receptors and transporters, and has been shown to inhibit hepatitis B virus replication [64,65]. This latter effect is in part achieved by stimulation of Ca^2+^ channels by Evans blue, resulting in reduced cytosolic levels of Ca^2+^. A similar mechanism might contribute to the anti-EBOV effect of Evans blue, as several processes in the EBOV life cycle, including fusion and budding, are affected by cytosolic Ca^2+^ concentrations [66-68]. UNC1999 is an inhibitor of the lysine methyltransferases enhancer of zeste homolog 1/2 (EZH1/2) [69]. It has primarily been studied as a potential anti-cancer drug because of its potential to alter the differential expression of host genes through epigenetic regulation [70]. Likewise, the mechanism through which UNC1999 inhibits EBOV replication might be that it counteracts the pro-viral manipulation of host factor pathways during EBOV infection [71]. Like UNC1999, benproperine, a clinically used antitussive drug, has also been evaluated as a potential anti-cancer drug. It has been shown to inhibit Actin-related protein 2/3 complex subunit 2, which plays a role in actin polymerization [72]. The transport of EBOV nucleocapsids to the cellular membrane prior to virion formation is dependent on actin polymerization, providing a potential explanation for the anti-EBOV mechanism of benproperine [73]. A final compound that showed highly selective inhibition of EBOV replication (SI >9), albeit with slightly lower potency (IC_50_ = 10.83), was doramapimod. This pyrazole-urea compound, originally developed for the treatment of inflammatory diseases, is a direct inhibitor of p38 mitogen activated protein (MAP) kinase [74]. This kinase is involved in the host cellular interferon type I response pathway and is indirectly inhibited by EBOV VP24 in certain cell types [75]. Furthermore, inhibitors of p38 MAP kinase have previously been shown to impair EBOV entry [76].

By screening >4,200 drug or drug-like compounds for their potential to inhibit EBOV, we identified several new *in vitro* inhibitors of EBOV replication. Although further validation of these compounds is needed, the use of a replication-competent virus-based assay ensures the direct biological relevance of the results shown here. Most compounds are active in the low micromolar range and display only limited cytotoxicity, making them good compounds to study EBOV replication and to potentially serve as a basis for the development of new therapeutics. Many top hits are also known or presumed to target different aspects of the viral life cycle, opening up the possibility for combination studies. Moreover, several of these compounds have favorable pharmacokinetic properties or have already been used as human therapeutics for other applications, making them valuable candidates for *in vivo* validation and potential further applications in the fight against EBOV.

## Acknowledgments

The authors wish to thank the Medicines for Malaria Venture and CD3 for providing compounds and Prof. S. Becker, Philipps-Universität, Marburg, Germany for providing the T7-3M-Luc-5M minigenome and EBOV support plasmids. B.V. was supported by a FWO SB grant for strategic basic research of the “Fonds Wetenschappelijk Onderzoek”/Research foundation Flanders (1S28617N). Part of this research work was performed using the ‘Caps-It’ research infrastructure (project ZW13-02) that was financially supported by the Hercules Foundation (FWO) and Rega Foundation, KU Leuven.

## Author contributions

This study was conceived by BV and PM. Experimental work was performed by BV and JSt. High-content imaging was performed by WC and JSc. Data analysis was performed by BV. AM, PC, PM and KV supplied reagents and materials. BV and PM drafted the manuscript. All authors read and approved the final version of the manuscript.

